# Modelling Medial Degenerative Features by Enzymatic Digestion to Evaluate Disease-Relevant Structure-Function Relationships in the Thoracic Aorta

**DOI:** 10.64898/2026.02.04.703920

**Authors:** Daniella Eliathamby, Lucas Ung, Hayley Yap, Malak Elbatarny, Maral Ouzounian, Michelle Bendeck, Michael A. Seidman, Craig A. Simmons, Jennifer C. Y. Chung

## Abstract

**Background:** Aortic microstructure-function relationships and the pathophysiology of how medial degeneration leads to aortic dissection remain poorly defined. We aimed to determine how degeneration of individual components of the extracellular matrix (ECM), namely elastin, collagen, and proteoglycans, influence biomechanical properties of aortic tissue through an improved, disease-motivated enzymatic digestion framework.

**Methods:** Porcine aortic tissue was sectioned into 200 µm thick samples in the media, and progressively digested with elastase or collagenase for selective degradation of these ECM components. Full thickness human aortic tissues were treated with chondroitinase, hyaluronidase, and heparinase to completely remove proteoglycans. Biomechanical characterization was performed using planar biaxial tensile testing, from which low- and high-strain modulus, transition-zone behaviour, strain-energy density, and energy loss were derived. Degree of elastin fiber degradation was analyzed using two photon excitation fluorescence imaging. Analysis of collagen degradation was performed using picrosirius red staining under brightfield and polarized light. Alcian blue staining was used to evaluate proteoglycan content.

**Results:** Induced fragmentation and disorganization of elastin fibers reduced low-strain load bearing capacity, evidenced by reduced low-strain modulus, strain-energy density, and transition zone stress, along with reduced energy loss. Targeted collagen disorganization similarly reduced strain-energy density and decreased strain at the onset of transition, consistent with premature collagen recruitment, and was accompanied by reductions in high strain modulus and energy loss with increasing collagen degradation. Proteoglycan removal decreased energy loss and was found to modulate low- and high-strain behaviour, including reduced strain-energy density and strain at onset of transition, and increased high strain modulus.

**Conclusions:** Through targeted modelling of ECM degenerative features on aortic tissue mechanics, we have identified distinct disease-associated biomechanical roles for major matrix constituents, with overlapping effects. These findings delineate mechanical consequences of component-specific matrix degeneration while underscoring the complex, multifactorial nature of structure-function relationships in aortic disease.

## Introduction

The extracellular matrix (ECM) plays a central role in the structural integrity and mechanical behavior of the aortic wall [1], [2]. In healthy aortic tissue, concentric elastic laminae embed an intricate combination of smooth muscle cells, collagen, and proteoglycans to form the medial lamellar unit, the building block of aortic tissue [3]. Elastin fibers provide passive elastic recoil and contribute to aortic compliance, while smooth muscle cells regulate active contractility and modulate effective compliance of aortic tissue [2]. In contrast, collagen protects the lamellar unit from overstretching in end-systole [2], [3] and proteoglycans maintain interlamellar adhesion and viscoelastic behaviour [3], [4]. Abnormal alterations to the medial lamellar unit result in disruption of normal tissue mechanics and possible predisposition of the aortic wall to aneurysm formation and subsequently risk of failure as seen in aortic dissection [1], [5]. Such alterations, collectively referred to as medial degeneration, can include elastin fiber loss and fragmentation, as seen in Marfan’s syndrome; collagen degradation, as seen in Ehlers-Danlos syndrome; and smooth muscle cell loss with accumulation and pooling of proteoglycans, as seen in sporadic or age-associated aneurysms [2], [3], [5]. However, while these individual degenerative features are prominent and commonly tied to specific aortopathies, medial degeneration is seldomly isolated to a single constituent. Rather, all degenerative features coexist to varying degrees, making it difficult to determine how each component mechanistically contributes to the weakening and increased risk of mechanical failure of the aortic wall. Understanding how tissue mechanics is influenced by specific ECM alterations is therefore critical to elucidating the pathways through which the medial degeneration observed in aortic aneurysms translates to the highly fatal events of aortic dissection and rupture.

The role of ECM alterations in aortic biomechanics can be studied by selective enzymatic digestion of ECM components. Roach et al. were the first to utilize this technique, and they found that removal of elastin resulted in an earlier transition to the stiffer, second linear phase of the arterial stress-strain response, whereas the characteristic nonlinear response was lost following removal of collagen [6]. This demonstration of elastin-dominated low strain behavior and collagen-dominated high strain behavior has been further validated in aortic tissue by several groups [7], [8], [9], [10], [11]. However, while these studies provided insight on the overall contribution of elastin and collagen on arterial vessel response, complete removal of ECM components does not reflect the medial degeneration seen in aortic aneurysms.

Alternatively, mild and progressive enzymatic digestions could be applied and their associated contribution to aortic biomechanics measured. Progressive elastin digestion models revealed unexpected softening in aortic stress-strain response, which contrasts the stiffening response found with complete digestion [10], [12], [13], [14]. Similar softening responses have also been reported in studies achieving graded collagen digestion, demonstrating progressively increasing tissue compliance as collagen is degraded up to its complete removal [15], [16]. However, histological examination of these progressive digestions revealed that the enzymes typically profoundly eliminated elastin or collagen at the edges of the aortic tissue with inward digestion towards the middle of the specimens as the digestion progressed. These studies consequently captured non-physiological content-function relationships, rather than the intended structure-function mechanisms. Therefore, to clarify the structure-function relationships associated with thoracic aortic aneurysms, a better disease-relevant model of ECM degenerative features is needed.

In addition, proteoglycans are increasingly recognised as key regulators of aortic biomechanics, yet they have been less frequently targeted by enzymatic digestion models compared to elastin and collagen, and the available studies report inconsistent effects. Beenakker et al. evaluated proteoglycan depleted porcine tissues by atomic force microscopy and found reduced stiffness following proteoglycan removal [9]. In contrast, Ghadie et al. observed a stiffening response by similar indentation testing, along with reduced residual stress as measured by opening angle [17]. Finally, Mattson et al. reported no significant change in overall stiffness after proteoglycan depletion, however, found an earlier transition point of the nonlinear stress-strain curve and reduced stress relaxation response of aortic tissues tested [18]. Collectively, these findings illustrate the diverse contributions of proteoglycans to aortic mechanics and thereby emphasize the need to further investigate their role in disease-relevant aortic biomechanics.

To this end, the aim for this study was to establish mechanistic insight into how individual ECM components – elastin, collagen, and proteoglycans – affect the biomechanical properties of the aortic wall through novel disease-relevant enzymatic degradation models of both porcine and human aortic tissues.

## Methods

### Elastin and Collagen Digestion

For both elastin and collagen digestion models, porcine aortic tissue was chosen due to its close structural and biomechanical similarity to that of the human aorta, and its availability for controlled, reproducible experimentation. Porcine hearts with full aortic specimens from pigs of unspecified age were obtained from a local slaughterhouse, and the ascending aorta was then isolated. Four to six 5 mm x 7 mm (circumferential x longitudinal) sections were taken along the outer curvature of each aorta for further processing (Supplemental Figure 1). To achieve uniform progressive degradation of these ECM components, thin sections of the media were obtained, allowing the enzymes access to the entirety of the aortic tissue. Briefly, full thickness sections were then cut down using a vibratome to achieve between four to six serial 200 µm thick sections of the medial layer from each 5 mm x 7 mm sample for subsequent enzymatic digestions. This sample thickness optimized mitigating the described edge effects from previous ex-vivo enzymatic digestions using intact tissues and facilitated tissue handling [12], [14].

For selective enzymatic digestion, 200 µm thick porcine aortic sections were treated with either elastase (2.5 U/mL, Millipore Sigma E1250) [12], [19] or purified collagenase (500 U/mL, Worthington LS005273) [10], [15] at 37°C. For each elastin digestion run, four serial sections were prepared, with one sample attached to a CellScale Biotester to assess planar biaxial mechanical properties following 0, 2, and 3 hours of digestion, and the remaining three sections incubated in the same solution and removed at the corresponding time points for imaging analysis. For each collagen digestion run, a total of seven serial sections across adjacent samples were prepared and treated similarly for digestion at 0, 1, 2, 3, 4, 5, and 6 hours. After removal from solution, all samples were rinsed with PBS and embedded in optical cutting temperature (OCT) medium for subsequent imaging analysis of ECM structural alterations. A total of ten (n=10) digestion runs were performed for each enzyme.

### Proteoglycan Digestion

Healthy aortic tissues demonstrate minimal tissue hysteresis [20], [21]. Therefore, normal porcine aortas were a poor model for studying the viscoelastic properties expected to be affected by proteoglycans. Therefore, for the proteoglycan degradation model, human aneurysmal aortic tissues, which our group previously demonstrated had increased viscoelastic hysteresis, were used [20], [21]. This study was approved by the research ethics board of the University Health Network, Toronto (16-6285, July 27, 2017), and all participants provided written informed consent. Briefly, whole ring ascending aortas from patients treated electively for aortic aneurysm were collected from the operating room and immediately placed on ice, with orientation marked by the surgeon. After clearing loose connective tissue from the adventitia, a custom 14 x 14 mm^2^ square cutter was used to collect two adjacent samples from each aorta from the anterior region, maintaining orientation. One sample was immediately embedded in OCT for evaluation of ECM prior to enzymatic treatment. The other sample was tested mechanically, followed by enzymatic digestion with 0.075 U/mL chondroitinase ABC (Sigma-Aldrich C3667), 15 U/mL hyaluronidase (Sigma-Aldrich H3506), and 0.75 U/mL heparinase (Sigma-Aldrich H3917) in 100 mM ammonium acetate buffer (pH 7.0) (Sigma-Aldrich A1542) for 48 hours at 37°C [17], [18]. Mechanical testing was then repeated after the digestion, and the sample rinsed in PBS and embedded in OCT medium for subsequent analysis.

### Mechanical Testing

Mechanical testing was adapted from previously published protocols [20], [21], [22], [23], [24]. Graphite powder was applied to all samples prior to testing to allow for optical strain tracking. For elastin and collagen models, 5 mm x 7 mm samples were trimmed to 4 mm squares prior to loading onto the Biotester (CellScale, Waterloo, ON, Canada) using tungsten biorakes. Samples were subjected to 10 preconditioning cycles at 25% rake-to-rake engineering strain, followed by multiple test cycles collected between 25-60% strain, with 4 test cycles obtained per strain level. Tissue thickness at each incubation time point was determined from serial samples removed for histological analysis using a high-magnification (12x) zoom lens (Navitar). For proteoglycan digestion, 14 x 14 mm^2^ square samples were mounted onto the Biotester using GoreTex sutures looped through each side 5 times. This allowed for the samples to be mounted and dismounted between 0-and 48-hour time points, as it was not possible to maintain the enzymatic solution in the Biotester water bath for a 48-hour duration. Samples were subjected to 10-20 preconditioning cycles at 30% rake-to-rake strain (with additional cycles to remove slack in suture lines), followed by 3 test cycles collected at both 30% and 35% strain. Tissue thickness was measured prior to 0-hour testing and after 48-hour testing. Engineering stress and engineering strain measured optically were used for all calculations.

Low strain modulus (LSM) was calculated as the secant modulus between the toe region up to the transition point, marking the first linear region of the aortic stress-strain curve. High strain modulus (HSM) was calculated as the tangent modulus at maximum achieved strain. These parameters capture modulus of elasticity under low versus high strain, respectively, and reflect aortic tissue stiffness. Transition zone strain (TZo strain) and stress (TZo stress) were defined as stress and strain values at the first break from linearity on the aortic stress strain curve (or transition point), determined through previously described change-point analysis [24]. Elastic energy per unit tissue volume (or strain energy density) was calculated as the area under the loading phase of the stress-strain curve up to the transition point. Finally, energy loss was calculated as area between loading and unloading stress-strain curves, normalized to area under the loading curve. This measures aortic tissue hysteresis and represents energy dissipated during cyclic loading. A schematic of the aortic stress-strain plot can be found in Supplemental Figure 2. Definitions and equations for biomechanical parameters derived from planar biaxial tensile testing data are summarized in Supplemental Table 1.

### Histologic Staining and Imaging

Multimodal histopathologic staining and imaging were performed to assess both primary and secondary effects of enzymatic digestion on ECM components. OCT-embedded tissue blocks from each digestion run were sectioned at 5 µm thickness for subsequent staining and imaging. For elastin, two-photon excitation fluorescence (TPEF) imaging was used to visualize and quantify elastin fiber structure (Nikon A1R MP Upright Multiphoton Microscope, 20x, 0.8 numerical aperture water immersion objective). Excitation wavelength was set to 810 nm and emission collected at 525 ± 25 nm [13], [19]. To measure elastin fragmentation, elastin fibers were traced using the ImageJ plug-in NeuronJ. Additionally, fiber waviness and organization were captured using the Directionality plug-in of ImageJ. Additional methodological detail is provided in Supplemental Materials.

For collagen, picrosirius red (PSR) staining was used to visualize collagen fibers (details provided in Supplemental Materials). Under brightfield, mean pixel intensity was measured as a quantitative indictor of collagen content and density, with higher intensity values corresponding to greater collagen content and more tightly packed fibers [25]. Mean pixel intensity was reported in arbitrary units (AU). PSR stained slides were also imaged under quantitative polarized light microscopy to evaluate collagen structure and organization (Abrio Imaging System, Cambridge Research and Instrumentation) [26]. The system measures birefringence of fibrillar collagen via the retardance of plane polarized light, and using a circular polarizer to detect orientation independent signal, provides an unbiased, quantitative measure of fibrillar collagen organization and density. [26], [27].

For proteoglycans visualization, alcian blue staining (pH 2.0) was used and imaged under brightfield microscopy (details provided in Supplemental Materials). Area fraction of proteoglycans was then calculated as area of blue pixels divided by total tissue area of slide [20]. Additionally, mean pixel intensity was measured, following methods outlined for collagen and PSR staining, in which higher intensity values correspond to greater relative concentration of proteoglycans within a given area [25]. Mean pixel intensity was reported in arbitrary units (AU).

### Statistics

All statistical analyses were performed using GraphPad Prism (v10.0.0). Data are presented as mean ± standard deviation. Repeated measures one-way analysis of variance (ANOVA) with Tukey’s post-hoc multiple comparisons test was used to assess differences in mechanical properties and histological outcomes across incubation time points for elastin and collagen digestion treatments. For proteoglycan digestion experiments, paired t-tests were used to compare measurements before and after enzymatic digestion. Statistical significance was defined as p<0.05.

## Results

### Elastin Degradation Modelling

Effects of the elastase treatment are displayed in Figure 1. Prior to elastase treatment (0-hour), elastin fibers were organized in concentric and continuous lamellae (Figure 1A), with average fragment lengths of 639.6 ± 141.9 µm (Figure 1B) and average waviness of 12.60 ± 3.49° (Figure 1C). All fibers were well-aligned with the circumferential axis (left to right in Figure 1A). After 2-hours of elastase exposure, discernable breaks and discontinuities in elastin architecture appeared throughout the full thickness of the sample, indicating progressive and uniform fiber fragmentation throughout the sample. Average fragment length after 2-hours significantly decreased from 0-hours (328.1 ± 79.29 µm, p=0.003), however average fiber waviness did not differ (16.32 ± 5.18°, p=0.35). After 3-hours exposure, significant alterations were observed in both average fiber length and waviness of fibers. Average fiber length after 3-hours (227.5 ± 69.92 µm) decreased compared to both 0-hour (p=0.0002) and 2-hour incubations (p=0.0073), and average waviness of fibers significantly increased from 0-hour (25.46 ± 9.91°, p=0.02). After 3-hours, fibers appeared overall shorter with less directional consistency, and greater interlamellar distance between adjacent fibers compared to 0-hour.

**Figure 1:**
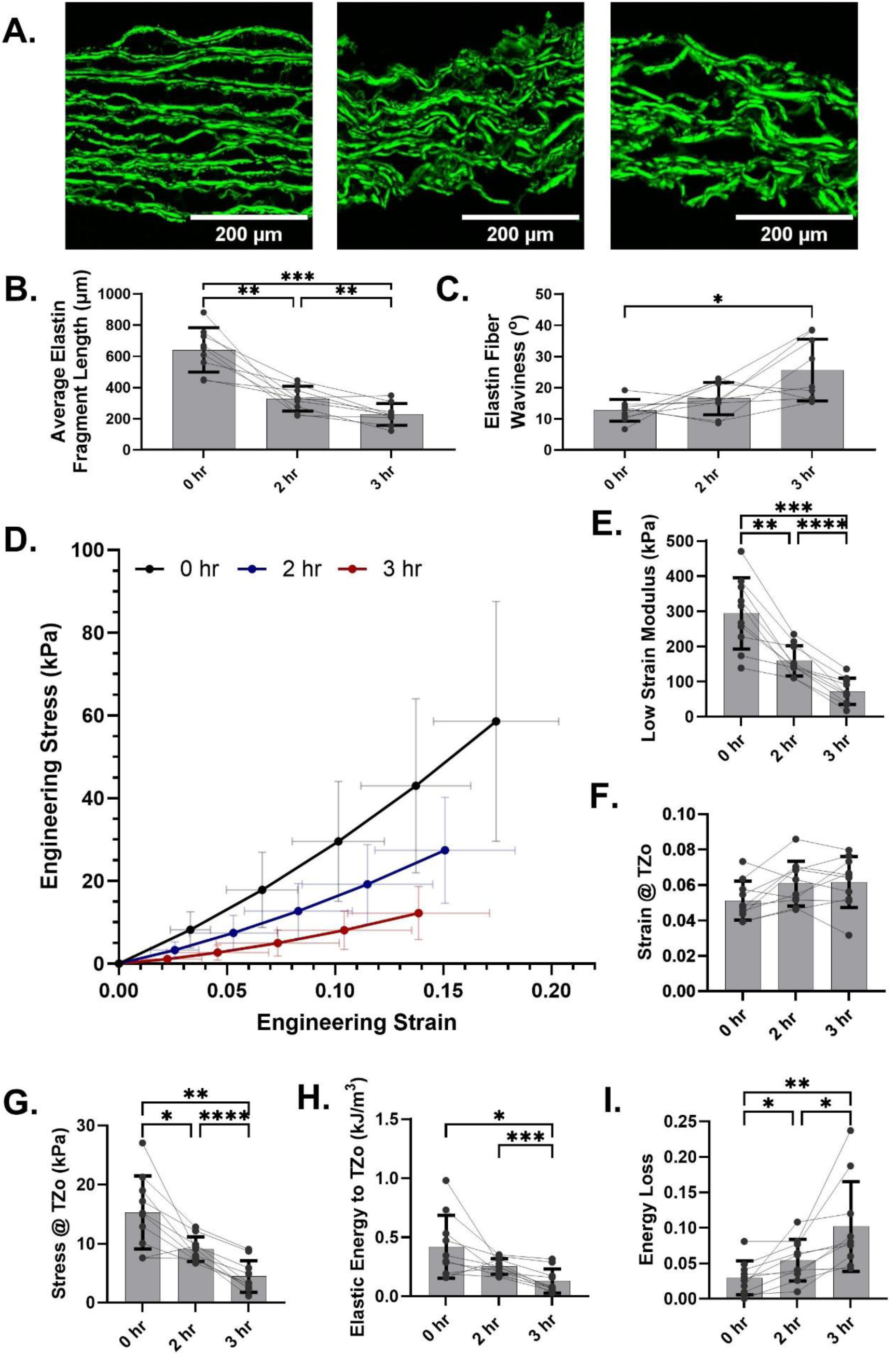
Effects of elastase treatment on elastin structure and mechanical properties. Two photon excitation fluorescence (TPEF) revealed significant alterations to the elastin network following 0-, 2-, and 3-hours of elastase treatment (A). Waviness of elastin fibers significantly increased after 3-hours incubation (B). Elastin fragmentation progressively increased from 0- to 3-hours (C). Average stress-strain curves revealed distinct softening behaviour following each treatment (D). Low strain modulus progressively and significantly decreased following treatments (E). No difference to transition zone strain was observed (F). However, transition zone stress decreased following 2- and 3-hour digestion (G). Elastic energy up to the transition zone decreased after 3-hours digestion (H). Energy loss increased after 3-hours (I). All comparisons were performed using one-way analysis of variance (ANOVA) with Tukey’s post-hoc multiple comparisons test (N=10). *p=0.01, **p=0.001, ***p=0.0001, ****p<0.001. Bars denoted mean ± standard deviation with individual paired points overlaid. Results are reported for the circumferential axis. Longitudinal data exhibited similar trends and are shown in Supplemental Figure 3.

Mechanical differences following elastase treatment are displayed in Figure 1D. Maximum optically tracked strain was matched from different rake-to-rake conditions tested, such that maximum achieved strain were similar amongst treatment times. Average loading stress-strain curves displayed a distinct softening behaviour from 0- to 3-hours of elastase incubation. This was reflected in significant changes to low strain modulus, with average low strain modulus at 0-hour (293.3 ± 101.3 kPa) greater than both 2-hour (158.7 ± 42.88 kPa, p=0.001) and 3-hour (72.21± 37.07 kPa, p=0.0002) incubations (Figure 1E). No significant differences in TZo strain were observed amongst treatments (Figure 1F), however, TZo stress decreased significantly from 0- to 2-hours (15.27 ± 6.189 kPa vs. 9.07 ± 2.08 kPa, p=0.02), 0- to 3-hours (4.42 ± 2.69 kPa, p=0.001), and 2- to 3-hours (p<0.0001) (Figure 1G). Elastic energy significantly decreased from 0- to 3-hours (0.13 ± 0.10 kJ/m^3^, p=0.0007) (Figure 1H). Energy loss significantly increased between 0- and 2-hour incubation (0.030 ± 0.024 vs. 0.054 ± 0.029, p=0.04), and 0 to 3 hours (0.102 ± 0.063, p=0.002) (Figure 1I). Due to the marked fragility of elastase-treated tissues, samples could not be sufficiently strained for calculation of high strain modulus.

Secondary alterations to collagen and proteoglycan networks from the applied elastase treatment are captured in Figure 2. Mean pixel intensity of collagen, measured by brightfield imaging of picrosirius red-stained sections, did not significantly differ between 0-, 2-, and 3-hours (39.46 ± 9.46 AU vs. 35.90 ± 10.14 vs. 40.26 ± 12.71 AU, ANOVA p=0.59), indicating similar overall content of collagen content between treatments (Figure 2A left side and 2B). However, retardance of collagen decreased significantly from 0- to 3-hours (12.91 ± 1.5 nm vs. 9.33 ± 2.78 nm, p=0.001), and between 2- and 3-hours (10.78 ± 2.93 nm, p=0.05) (Figure 2A left side and 2C). Area fraction of proteoglycan decreased significantly between 0- and 3- hours (0-hr: 22.69 ± 4.87% vs. 3-hr: 16.83 ± 3.49%, p=0.03) and 2- and 3-hours (2-hr: 20.52 ± 3.36%, p=0.01) of digestion (Figure 2D and 2E). Similarly, alcian blue mean pixel intensity decreased between 0-and 2-hours (24.99 ± 5.38 AU vs. 21.21 ± 5.54 AU, p=0.007), but not between 0- and 3-hours of incubation (26.08 ± 8.19 AU, p=0.85) (Figure 2D and 2F). Instead, mean pixel intensity was found to increase from 2- to 3-hours (p=0.03), which suggested pooling of remaining proteoglycans with degradation of elastin.

**Figure 2:**
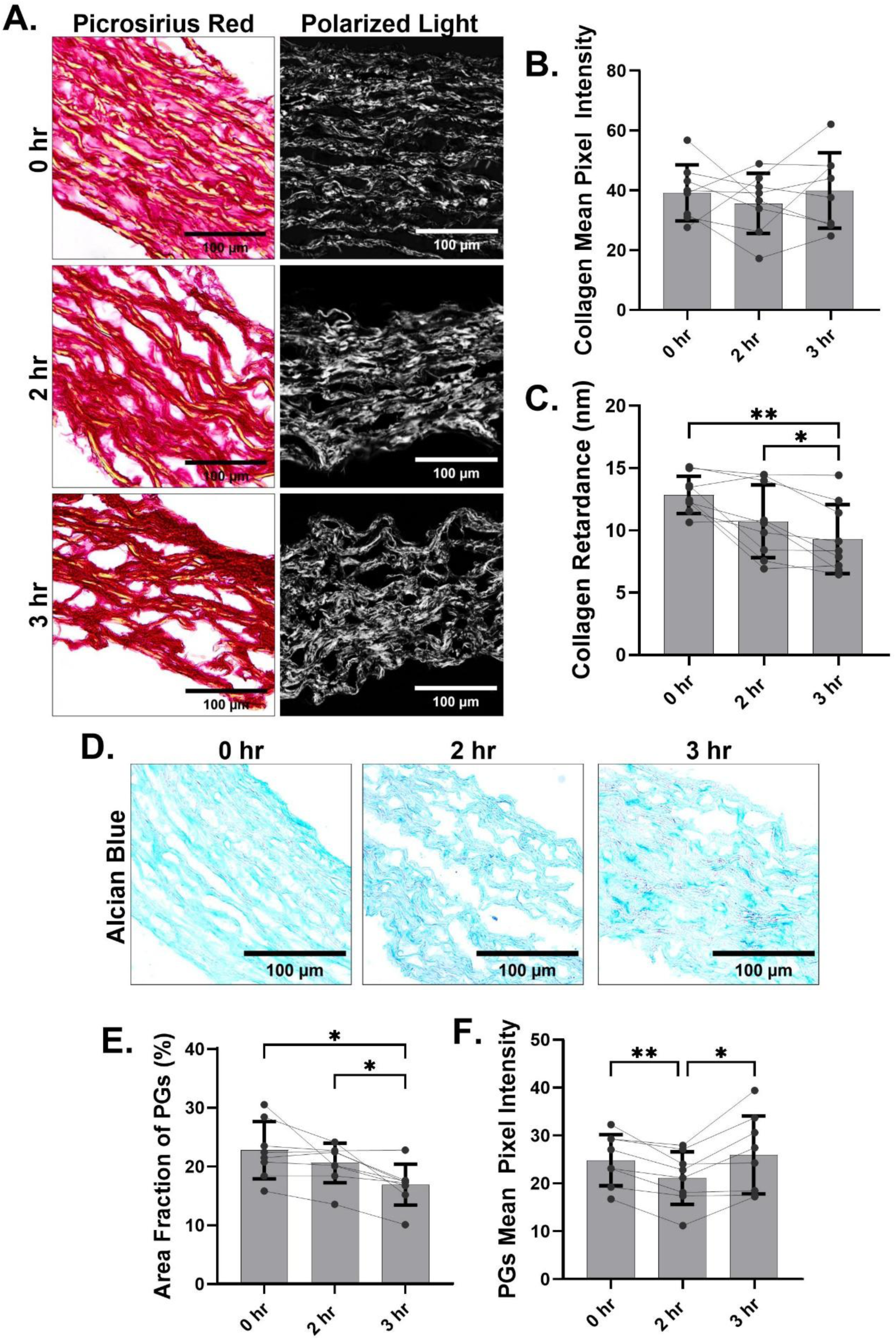
Secondary alterations to collagen and proteoglycan networks following elastase treatment. Picrosirius red stained slides under brightfield (left) and polarized light (right) for 0-, 2-, and 3-hour elastase treatments demonstrated simultaneous secondary alterations to collagen as a result of elastin degradation (A). Mean pixel intensity did not significantly differ between treatments (B). However, collagen retardance significantly decreased following 3-hour treatment (C). Alcian blue stained slides under brightfield demonstrated secondary alterations to the proteoglycan network after elastase treatment (D). Area fraction of proteoglycans progressively decreased following elastase digestion (E). Mean pixel intensity decreased from 0- to 2-hours, however, increased after 3-hours compared to 2-hours incubation (F). All comparisons were performed using one-way analysis of variance (ANOVA) with Tukey’s post-hoc multiple comparisons test (N=10). *p=0.01, **p=0.001, ***p=0.0001, ****p<0.001. Bars denote mean ± standard deviation with individual paired points overlaid.

### Collagen Degradation Modelling

Effects on the collagen network following collagenase treatment are shown in Figure 3. Prior to treatment (0 hr), collagen mostly appeared dark red in picrosirius red-stained sections (Figure 3A left). Mean pixel intensity (60.05 ± 8.902 AU) and retardance (11.62 ± 2.45 nm) were highest pre-treatment, compared to all other treatment times, indicating more dense and well-organized fibers prior to digestion. After 3 hours digestion, significantly lower mean pixel intensity (19.75 ± 4.90 AU, p=0.004) and retardance (4.51 ± 1.50 nm, p=0.03) were observed. Relative to 0-hour, concentrated red staining was diminished, with larger apparent gaps between lamellae. This pattern continued up to 6-hours with decreased pixel intensity (15.17 ± 3.28; p=0.002 relative to 0-hour), decreased retardance (1.76 ± 0.42 nm; p=0.001), and large interlamellar voids observed (Figure 3A-3C).

**Figure 3:**
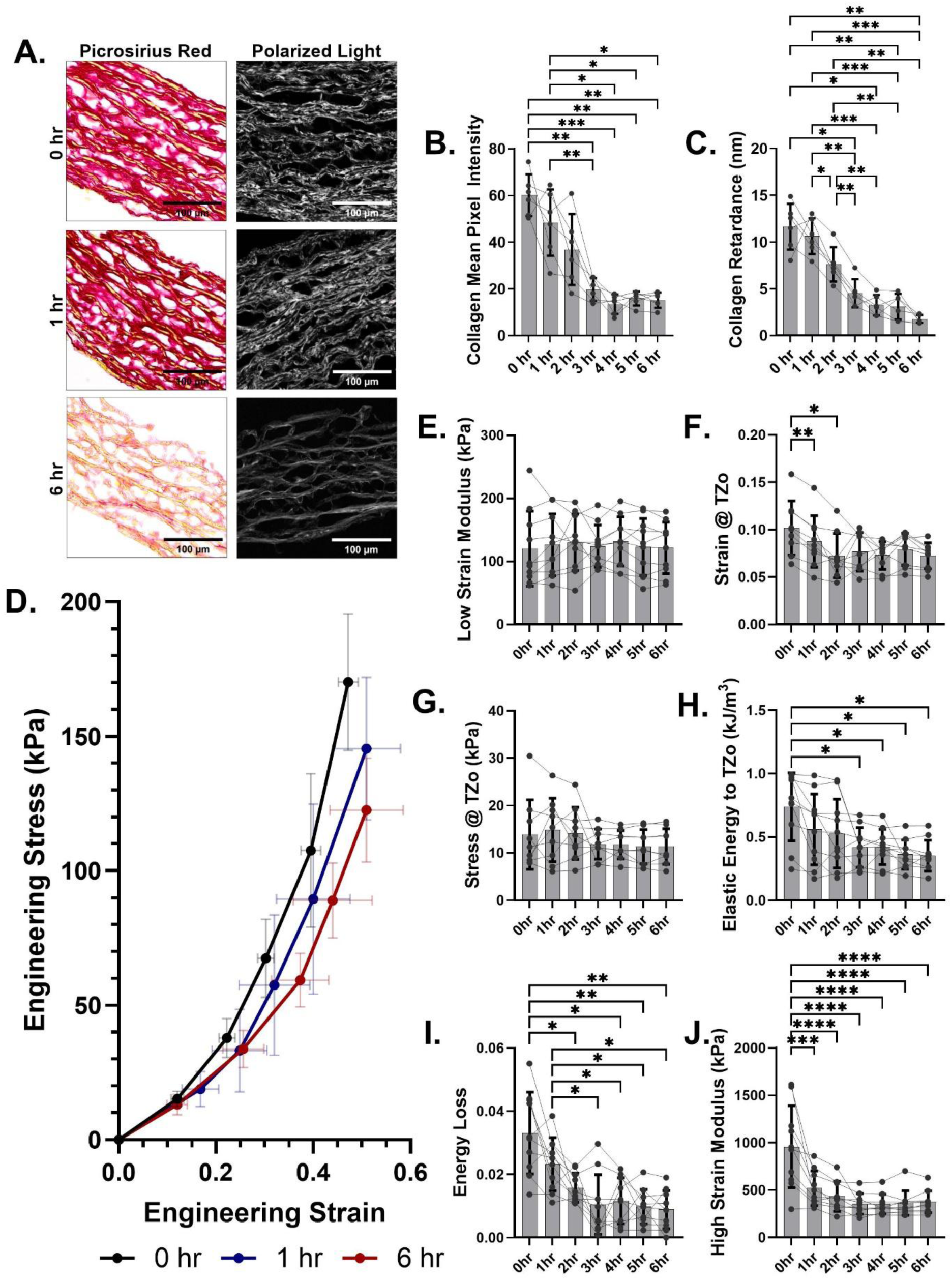
Effects of collagenase treatment on collagen organization and mechanical properties. Picrosirius red stain under brightfield (left) and polarized light (right) confirmed alterations to collagen at 0-, 1-, and 6-hour timepoints (A). Mean pixel intensity significantly dropped after applied collagenase treatment, indicating removal of collagen (B). Similar trends were also observed in retardance, indicating increased disorganization of remaining collagen (C). Average stress-strain plots revealed similar toe-region behaviour but notable softening at higher strain compared to 0-hr (D). Low strain modulus was relatively stable despite removal and disorganization of collagen (E). Decreased strain at the transition zone (TZo) was observed after mild treatment (F). No differences to stress at the transition zone were observed (G). A drop in elastic energy up to the transition zone was evident following 3-hours of digestion (H). Energy loss and high strain modulus exhibited significant decreases as collagenase treatment was applied (I and J). All comparisons were performed using one-way analysis of variance (ANOVA) with Tukey’s post-hoc multiple comparisons test (N=6). *p=0.01. **p=0.001. ***p=0.0001. ****p<0.001. Bars denote mean ± standard deviation with individual paired points overlaid. Representative picrosirius red and polarized light images from all tested timepoints can be found in Supplemental Figure 4, and all average stress-strain plots in Supplemental Figure 5. Results are reported for the circumferential axis. Longitudinal data exhibited similar trends and are shown in Supplemental Figure 6.

Changes to mechanical properties following collagen alterations are displayed in Figure 3D. Low strain-regions of the aortic stress-strain curves were on average similar between treatments, with low strain modulus consistent across all treatment times (p_ANOVA_=0.99) (Figure 3E). Compared to 0-hour, strain at the transition zone significantly decreased following 1- (0.10 ± 0.03 vs. 0.09 ± 0.03, p=0.001) and 2-hours of digestion (0.10 ± 0.03 vs. 0.07 ± 0.02, p=0.01) (Figure 3I). No significant changes occurred to stress at the transition zone (p_ANOVA_=0.28) (Figure 3J). Elastic energy significantly decreased following 3-hour digestion (0.42 ± 0.16 kJ/m^3^, p=0.02), after which it plateaued. A significant decrease in energy loss was observed between 0 and 3-hours of incubation (0.033 ± 0.013 vs. 0.010 ± 0.009, p<0.0001), with no further changes with longer digestions. After 1 hour digestion, there was a significant drop in high strain modulus compared to 0-hour (956.4 ± 432.3 kPa vs. 518.9 ± 182.5 kPa, p=0.0002) and remained relatively unchanged thereafter.

Figure 4 summarizes associated alterations to elastin and proteoglycan structure and content following collagenase treatment. Waviness of elastin fibers were significantly altered as a result of collagen digestion (p_ANOVA_=0.002). Waviness of fibers significantly increased between 0- and 1-hour digestion (6.64 ± 1.66° vs. 9.64 ± 1.79°, p=0.009). Between 4- and 6-hours, further notable and significant increase in waviness was observed (p=0.03) (Figure 4A and 4B). Marginally significant differences in average elastin fragment length were observed across treatments (p_ANOVA_=0.06), with elastin fibers after 6 hours of digestion shorter than those at 0-(471.9 ± 102.2 µm vs. 311.6 ± 59.97 µm, p=0.06) and 1-hour digestion timepoints (454.6 ± 88.06 µm vs. 311.6 ± 59.97 µm, p=0.08) (Figure 4A and 4C). Significant alterations were also observed in proteoglycan network following collagenase treatment (Figure 4D). A progressive drop in area fraction of proteoglycans was observed between 0- and 6-hours (p_ANOVA_<0.0001). Compared to 0-hours (30.54 ± 4.51%), the 6-hours digestion (17.05 ± 2.80%, p=0.004) demonstrated significantly less proteoglycan content. Intensity of alcian blue stain by mean pixel intensity was also affected by the collagenase digestion. Overall, there was a steady decrease in mean pixel intensity from 0- (59.90 ± 10.51) to 6-hours (38.19 ± 6.05) (p_ANOVA_=0.0007), which indicated no redistribution of remaining proteoglycans with collagen degradation in contrast to elastin degradation (Figure 4F).

**Figure 4:**
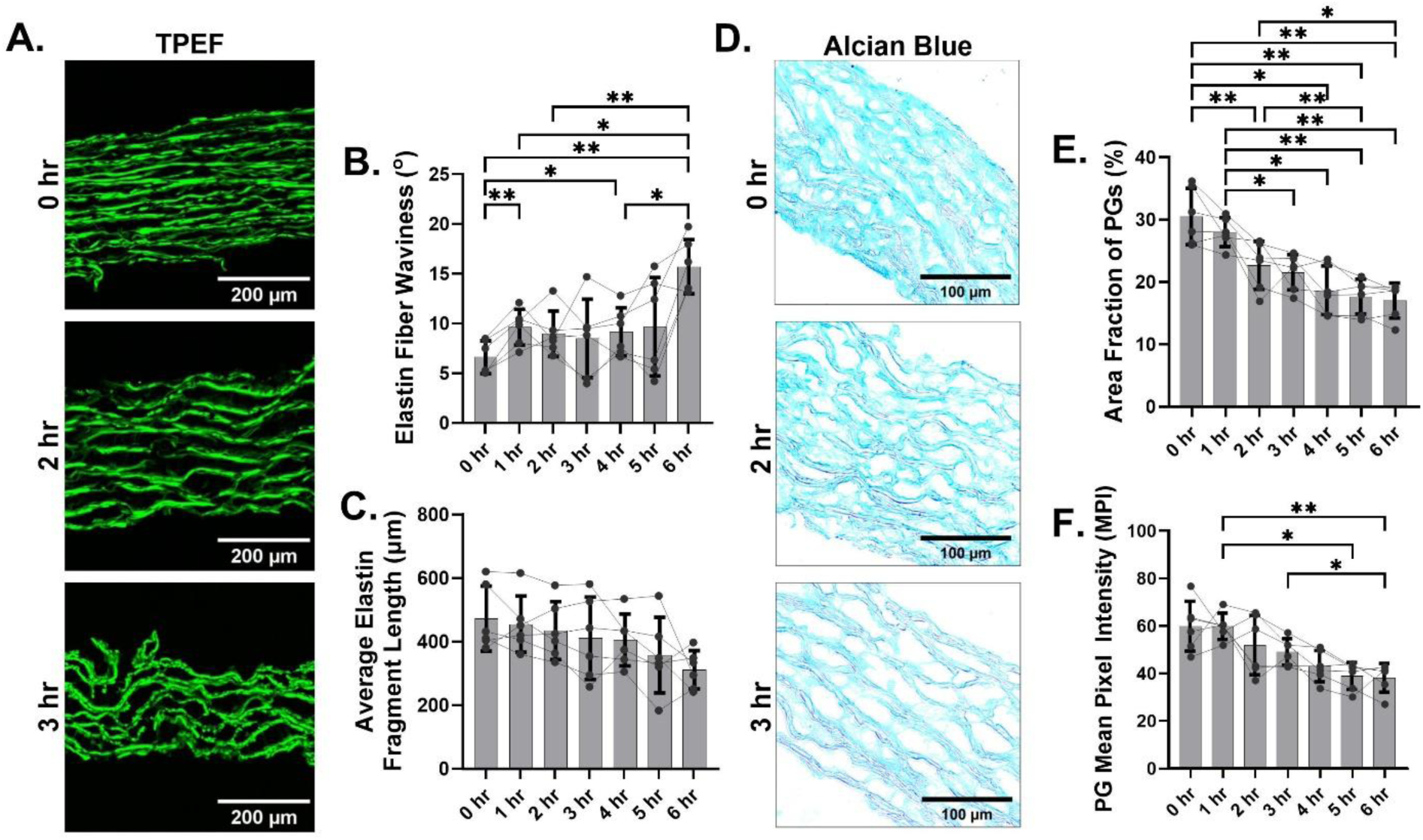
Secondary alterations to elastin and proteoglycan networks following collagenase treatment. Two photon excitation fluorescence (TPEF) imaging of elastin showed consequential effects to the elastin network following removal and degradation of collagen after 0-, 3-, and 6-hours of treatment (A). Waviness and dispersion of elastin fibers were significantly increased at 1-, 4-, and 6-hrs digestion of collagen, compared to 0-hr. (B). However, no difference in average length of elastin fibers was observed (C). Effects of collagenase treatment of proteoglycans are shown through alcian blue staining imagined under brightfield (D). Area fraction of proteoglycans was significantly decreased as a result of collagen digestion (E). In contrast, fewer effects were observed to mean pixel intensity (F). All comparisons were performed using one-way analysis of variance (ANOVA) with Tukey’s post-hoc multiple comparisons test (N=6). *p=0.01. **p=0.001. ***p=0.0001. ****p<0.001. Bars denote mean ± standard deviation with individual paired points overlaid.

### Proteoglycan Degradation Modelling

Effects on the proteoglycan network from applied enzymatic treatment are displayed in Figure 5. Following the 48-hour incubation, most proteoglycans were removed from the ECM. Area fraction of proteoglycans decreased significantly from pre-treatment (0-hr: 37.61 ± 8.71%) to post-treatment (48-hr: 9.51 ± 4.01%) (p<0.0001) (Figure 5A and 5B). Compared with the pre-treatment proteoglycans, the mean pixel intensity of those remaining post-treatment was significantly reduced (92.35 ± 16.43 vs. 33.11 ± 11.05, p<0.0001) (Figure 5A and 5C).

**Figure 5:**
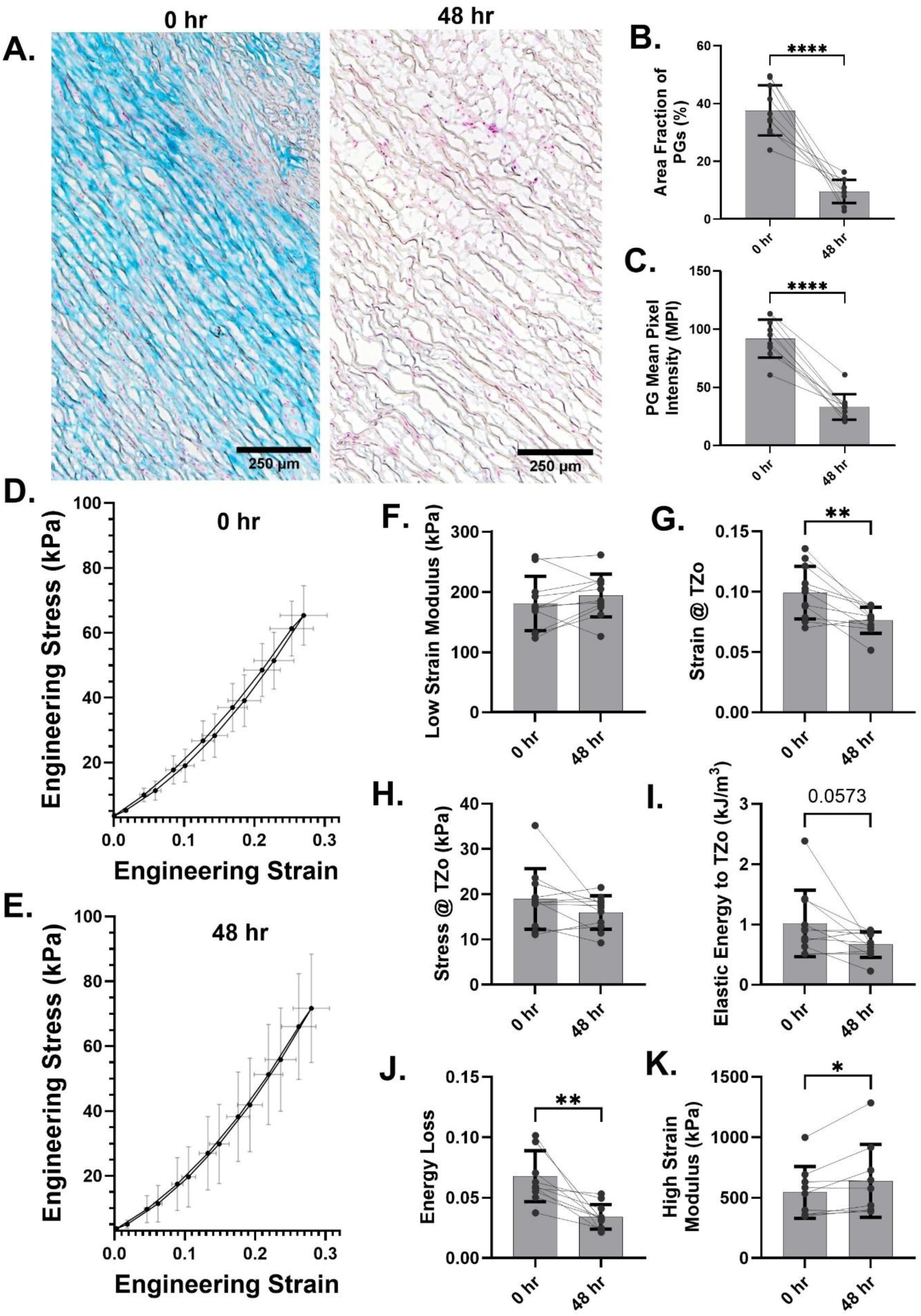
Histological and biomechanical effects following proteoglycan removal by enzymatic digestion. Digestion was confirmed by alcian blue staining (A). Area fraction of proteoglycans and mean pixel intensity of alcian blue staining were significantly reduced after digestion, indicating successful removal of proteoglycans from aortic tissues (Figure B and C). Pre-treatment, tissue hysteresis on average aortic stress-strain curve was evident (D). Post-treatment, minimal hysteresis was observed (E). No consistent trend to alterations in low strain modulus were reported (F). Strain at the transition zone was significantly decreased (G), but no difference to stress at the transition zone was observed following removal of proteoglycans (H). Energy stored up to the transition one was decreased following removal of proteoglycans with marginal significance (I). Energy loss was significantly decreased following digestion (J). High strain modulus was found to be significantly increased with removal of proteoglycans (K). All comparisons were performed using paired t-test (N=10). *p=0.01. **p=0.001. ***p=0.0001. ****p<0.001. Bars denote mean ± standard deviation with individual paired points overlaid. Results are reported for the circumferential axis. Longitudinal data exhibited similar trends and are shown in Supplemental Figure 7.

Biomechanical effects from removal of proteoglycans are also displayed in Figure 5. Tissue hysteresis was notably reduced, as displayed in average stress-strain plots (Figure 5D and 5E). No consistent changes in low strain modulus were found after 48-hour digestion (180.4 ± 45.20 kPa vs. 193.6 ± 35.63 kPa, p=0.21) (Figure 5F). Strain at the transition zone decreased following digestion (0.10 ± 0.02 vs. 0.08 ± 0.01, p=0.008) (Figure 5G). No changes to stress at the transition zone were observed (18.98 ± 6.72 kPa vs. 15.97 ± 3.70 kPa, p=0.14) (Figure 5H). There was a marginally significant decrease in elastic energy up to the transition zone after treatment (1.02 ± 0.55 kJ/m^3^ vs. 0.67 ± 0.21 kJ/m^3^, p=0.06) (Figure 5I). Energy loss decreased following removal of proteoglycans (0.068 ± 0.021 vs. 0.034 ± 0.010, p=0.001) (Figure 5J). High strain modulus marginally but significantly increased after removal of proteoglycans (554.6 ± 214.3 kPa vs. 640.0 ± 302.1 kPa, p=0.03) (Figure 5K).

Secondary effects to elastin and collagen structure and organization as a result of proteoglycan digestion are displayed in Figure 6. No consistent changes to elastin fiber waviness were observed (0-hour: 13.36 ± 4.72° vs. 48-hour: 11.95 ± 3.37°, p=0.51) (Figure 6A and 6B). Average elastin fiber length was not different before (486.1 ± 101.3 µm) and after (491.6 ± 103.0 µm) treatment (p=0.73) (Figure 6A and 6C). However, from TPEF imaging, collapse of interlamellar units was observed (Figure 6A). For collagen, prior to enzymatic digestion of proteoglycans, pink-red regions were observed under brightfield, indicating the presence of both thicker, tightly packed fibers (red) and thinner, less densely packed fibers (pink) (Figure 6D). After digestion, pink regions appeared reduced, with only intense red regions visualized around yellow-stained structures, as shown by increased mean pixel intensity (0-hr: 37.35 ± 10.87 vs. 48-hour: 46.37 ± 7.70, p=0.05). Additionally, retardance (Figure 6E) decreased post-digestion, although with marginal significance (0-hour: 15.97 ± 4.08 nm vs. 48-hour: 12.93 ± 2.27 nm, p=0.06) (Figure 6F).

**Figure 6:**
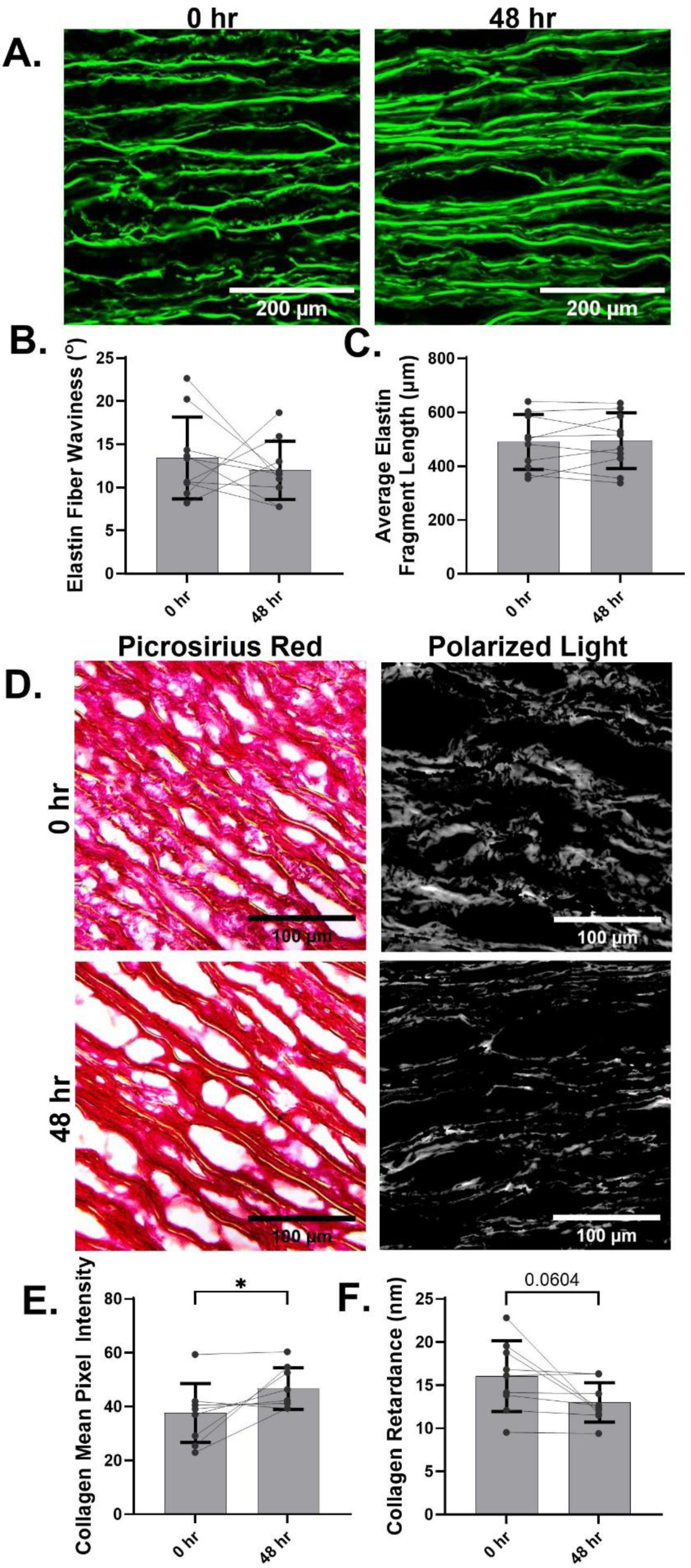
Secondary effects to elastin and collagen following removal of proteoglycans by enzymatic digestion. Two photon excitation fluorescence imaging display subtle but nonsignificant differences to the elastin network (A). Fiber waviness did appear to be altered following removal of proteoglycans; however, trends were inconsistent (B). Average fiber length of elastin fibers was similar pre- and post-digestion (C). Significant alterations were observed to collagen following proteoglycan digestion, in both brightfield and polarized light (D). Mean pixel intensity was significantly increased after digestion (E), and retardance was significantly decreased (F). All comparisons were performed using paired t-tests (N=10. *p=0.01. **p=0.001. ***p=0.0001. ****p<0.001. Bars denote mean ± standard deviation with individual paired points overlaid.

## Discussion

Through targeted digestion models, we evaluated the role of major ECM constituents on aortic tissue biomechanics. The novel use of thin layers of aortic media in these digestions models allowed for the study of controlled elastin fragmentation and disorganization, as well as collagen disorganization. Furthermore, the unique inclusion of human aneurysmal aortas in a proteoglycan digestion model allowed for study of aortic viscoelastic properties, which is increasingly recognized for its importance. Additionally, we considered the secondary effects of our digestion models on the other ECM constituents, providing insight into the interdependence of these networks on aortic biomechanics. In particular, these components were found to have a coordinated effect on the under-studied viscoelastic properties of the aortic wall.

Induced fragmentation and disorganization of elastin fibers was shown to reduce low strain modulus, as expected. This finding is in line with previous studies from Chow et al. and Lacolley et al., in which the degraded and removed elastin in whole arterial tissues exhibited greater distensibility compared to normal [12], [28]. Additionally, the transition to high-strain regions were found to occur at a lower stress level. This reduced stress at the onset of the transition zone reflects overall diminished load bearing capacity of the degraded elastin fibers. This is further supported by the reduced elastic energy up to the transition zone, capturing the reduced potential energy of degraded elastin fibers within aortic tissue.

Minor alterations to collagen fiber structure after 1-hour collagenase treatment, resulted in pronounced reduction in high strain modulus, reflecting the established role of collagen as the primary load-bearing component at high strain [6] [29], [30]. Cross-links restrain sliding between tropocollagen molecules (the structural unit of collagen), enhancing resistance to deformation [31], [32]. We suspect that targeted cleavage of tropocollagen by purified collagenase treatment created local fibril breaks, enabling molecular sliding and compromising network-level load sharing. Overall, these findings suggest that the increased, but structurally impaired collagen as seen in aneurysmal tissue may be mechanically abnormal and, rather than reinforce the vessel wall, paradoxically reduce high-strain modulus [33]. This is furthered supported by Pichamuthu et al. who found similar collagen content but significantly lower maximal tangential stiffness and tensile strength of aneurysm tissue from patients with tricuspid aortic valves compared to those with bicuspid valves, suggesting collagen structure and organization more than content dictate high strain behavior [34].

Longer collagen digestion treatments were shown to be associated with changes in the transition zone between the low- and high-strain regions of the aortic stress strain response, with decreased elastic energy up to the transition zone. This suggests a premature tradeoff from elastin- to collagen-dominated engagement following degradation to collagen, possibly related to the increased disorganization of elastin fibers observed alongside collagen digestion. However, while the elastic energy up to the transition zone primarily reflects the contribution of elastin, collagen also deforms elastically at low strains, associated with fibril uncrimping and straightening, [30], [31], [35]. Collagenase is suspected to alter collagen fiber crimp indirectly, as cleavage along the fiber reduces connectivity and therefore the prestress that maintains the crimped state when unloaded [36]. Therefore, our finding of reduced elastic energy and transition zone strain can be indicative of the uncrimping of collagen fibrils, which positions collagen for premature engagement independent of elastin architecture. These suspected alterations to collagen organization are further supported by imaging, where reduced intensity under brightfield and retardance under polarized light indicate less densely packed fibers, as expected with altered crimp and interconnectivity.

The most interesting findings of this study, however, are related to how the ECM components interacted to influence aortic wall viscoelasticity, which we measured through energy loss. Proteoglycans have been previously linked to viscoelastic behavior [3], [18], [37]. This is due to negatively charged glycosaminoglycan side chains that attract intra- and extrafibrillar water, acting as molecular lubricants that facilitate fibril sliding during tensile deformation [37]. This, coupled with the osmotic swelling from water recruitment, contribute to the energy dissipation capacity of proteoglycans, and thereby viscoelastic response [18]. The reduction in energy loss following proteoglycan digestion observed in our study provided further evidence of this effect. However, we found that alterations to elastin fibers also affected aortic viscoelasticity. Specifically, elastin fragmentation and disorganization resulted in a significant increase in energy loss. This finding supports recent mouse data, where the aortas from *Eln*-/-, *Lox*-/- and *Fbln*-/-mice (lacking elastin and/or proper assembly of elastin) exhibited increased dissipated energy compared to wild-type controls [38]. Previous in vitro studies of arterial elastin described the interaction of intrafibrillar water with elastin molecules, altering their stiffness and extensibility [39], [40]. In this context, fragmented and ill-assembled elastin may impair their water retention capacity. While proteoglycan content diminished in our elastin digestion model, proteoglycan aggregation, pooling and interlamellar spacing increased (Figure 2D), explaining increased tissue hysteresis [3], [38], [41]. These findings provide mechanistic insights into the correlation we previously reported between increasing energy loss and decreased levels of elastin in human aneurysmal aortic tissue [20].

Collagen digestion on the other hand resulted in a sharp decrease in energy loss. Similarly, studies in cartilage have reported reduced stress relaxation and loss moduli following collagen removal, further emphasizing a link between collagen and viscoelastic behavior of these soft tissues [42]. These enzymatic digestion studies of collagen in cartilage resulted in secondary and unintended removal of proteoglycans [43], [44]. In the present study, a similar co-removal of collagen and proteoglycans was observed, potentially reducing frictional interactions (and associated viscous energy losses) between collagen fibrils and the surrounding matrix [45]. The inverse may be reasonably hypothesized, that the increased deposition of collagen seen in medial degeneration increases proteoglycan recruitment and may contribute to the greater levels of energy loss demonstrated by thoracic aortic aneurysms.

Interaction between collagen and proteoglycans was also evident in our proteoglycan digestion model where removal of proteoglycans significantly reduced strain at the transition zone, while increasing high strain modulus. Findings of premature transition to high strain behavior were also reported by Mattson et al., and can be linked to the altered state of collagen structure as a result of prestresses removed from the fibrils after proteoglycan depletion [18]. This allows for the uncrimping of fibrils at lower applied strain, causing earlier engagement of collagen. However, contrary to our study, Mattson et al. did not find any significant differences to high strain modulus.

Understanding how ECM components interact to influence aortic viscoelasticity is significant because greater viscoelasticity has been implicated in increased risk of aortic dissection. Elevated energy loss correlates with weakening of the aortic wall (decreased delamination strength and ultimate tensile strength) and is seen in cases of aortic dissection [20], [46]. The proteoglycan pooling that we show may be contributed to by the elastin fragmentation and collagen deposition seen in medial degeneration which elevates Donnan pressures and localized swelling stresses. In turn these stresses reduce mechanical coupling between lamellar units and increase interlamellar shear stress. We have previously shown that once a critical amount of energy loss is reached, the adhesive strength between layers of the aortic media plummets resulting in a weakened aorta that would be prone to aortic dissection [20].

## Limitations

Our study is limited by the use of thinly sliced 200 µm medial sections of aortic tissue for enzymatic treatment. Inevitably, this alters the overall mechanical behavior compared to full thickness aortic tissue. However, this approach was necessary to ensure consistent and uniform digestion throughout the full thickness of the tested samples, addressing the edge effect limitations observed in previous enzymatic modeling studies. Moreover, as the objective of this work was to identify structural and mechanical trends rather than absolute property values, this limitation does not detract from the interpretation of our findings.

Our study is also limited by the sensitive nature of tissue following enzymatic digestion, which restricted the application of higher strain levels across all treatment conditions. This limitation is particularly relevant for the calculation of high strain modulus, which in the present study was derived from the tangent at maximum achieved strain rather than maximum strain prior to the yield point. Additionally, this limitation precluded combined enzymatic approaches targeting multiple ECM constituents in a single digestion, as samples were marginally manipulable at high degradation levels targeting a single constituent. Nonetheless, as our goal was to identify isolated disease associated structure-function relationships within the physiologically relevant range, the captured high-strain modulus and single target approach were considered appropriate. Future work will continue to explore both higher strain application and simultaneous digestion of ECM constituents using milder and shorter protocols to better capture the combined effects of ECM degradation on overall mechanical behavior.

Additionally, despite an extensive characterization of secondary effects following the digestion of each ECM component, our study was limited by its in vitro design, which cannot recapitulate the dynamic remodeling of living tissues over time in response to matrix degradation. Nevertheless, our goal was to better understand structure-function relationships relevant to in vivo disease processes. As such, future work will shift towards assessment of human aortic tissue, in which insights from these in vitro models can still be applied to investigate the highly complex and heterogeneous nature of human aortic tissue.

Finally, the exclusion of smooth muscle assessment represented a key limitation of this study. Vascular smooth muscle cells are widely considered to contribute primarily to active mechanical behavior, with minimal impact on passive mechanical properties of large elastic arteries [47]. Moreover, the viability of smooth muscle cells in resected aortic specimens is unclear, and potential cell death or phenotypic change ex vivo could bias mechanical measurements.

## Conclusions

This study identified distinct disease-associated biomechanical roles for major matrix constituents, advancing our understanding of how medial degeneration impacts aortic tissue integrity. Here, we found induced fragmentation and degradation of elastin fibers impaired elastin engagement within lower physiological strain ranges and reduced elastic recoil, resulting in a more compliant but mechanically less efficient tissue response, evidenced by increased greater energy loss during cyclic loading. Additionally, degradation of collagen led to a significant reduction to stiffness at higher strains, independent of the severity of alterations, suggesting that the structurally impaired collagen deposited alongside aneurysm formation may paradoxically reduce vessel compliance. Finally, removal of proteoglycans confirmed their role in maintaining viscoelastic behavior of aortic tissue but revealed their influence on both low- and high-strain mechanical properties, highlighting their contribution to overall tissue integrity.

## References

[1] P. Rai, L. Robinson, H. A. Davies, R. Akhtar, M. Field, and J. Madine, “Is There Enough Evidence to Support the Role of Glycosaminoglycans and Proteoglycans in Thoracic Aortic Aneurysm and Dissection?—A Systematic Review,” IJMS, vol. 23, no. 16, p. 9200, Aug. 2022, doi: 10.3390/ijms23169200.

[2] S. Jana, M. Hu, M. Shen, and Z. Kassiri, “Extracellular matrix, regional heterogeneity of the aorta, and aortic aneurysm,” Exp Mol Med, vol. 51, no. 12, pp. 1–15, Dec. 2019, doi: 10.1038/s12276-019-0286-3.

[3] J. D. Humphrey, “Possible Mechanical Roles of Glycosaminoglycans in Thoracic Aortic Dissection and Associations with Dysregulated Transforming Growth Factor-β,” J Vasc Res, vol. 50, no. 1, pp. 1–10, 2013, doi: 10.1159/000342436.

[4] Y. H. Shen, H. S. Lu, S. A. LeMaire, and A. Daugherty, “Unfolding the Story of Proteoglycan Accumulation in Thoracic Aortic Aneurysm and Dissection,” ATVB, vol. 39, no. 10, pp. 1899–1901, Oct. 2019, doi: 10.1161/ATVBAHA.119.313279.

[5] D. Costa, M. Andreucci, N. Ielapi, G. Accarino, U. M. Bracale, and R. Serra, “Aortic wall degeneration in aortic aneurysms. Pathological and molecular insights,” Annals of Vascular Surgery, p. S0890509625006740, Oct. 2025, doi: 10.1016/j.avsg.2025.10.002.

[6] M. R. Roach and A. C. Burton, “THE REASON FOR THE SHAPE OF THE DISTENSIBILITY CURVES OF ARTERIES,” Can. J. Biochem. Physiol., vol. 35, no. 8, pp. 681–690, Aug. 1957, doi: 10.1139/o57-080.

[7] A. Pukaluk, G. Sommer, and G. A. Holzapfel, “Multimodal experimental studies of the passive mechanical behavior of human aortas: Current approaches and future directions,” Acta Biomaterialia, vol. 178, pp. 1–12, Apr. 2024, doi: 10.1016/j.actbio.2024.02.026.

[8] N. Gundiah, A. R. Babu, and L. A. Pruitt, “Effects of elastase and collagenase on the nonlinearity and anisotropy of porcine aorta,” Physiol. Meas., vol. 34, no. 12, pp. 1657–1673, Dec. 2013, doi: 10.1088/0967-3334/34/12/1657.

[9] J.-W. M. Beenakker, B. A. Ashcroft, J. H. N. Lindeman, and T. H. Oosterkamp, “Mechanical Properties of the Extracellular Matrix of the Aorta Studied by Enzymatic Treatments,” Biophysical Journal, vol. 102, no. 8, pp. 1731–1737, Apr. 2012, doi: 10.1016/j.bpj.2012.03.041.

[10] A. J. Schriefl, T. Schmidt, D. Balzani, G. Sommer, and G. A. Holzapfel, “Selective enzymatic removal of elastin and collagen from human abdominal aortas: Uniaxial mechanical response and constitutive modeling,” Acta Biomaterialia, vol. 17, pp. 125–136, Apr. 2015, doi: 10.1016/j.actbio.2015.01.003.

[11] M. Kobielarz, A. Chwiłkowska, A. Turek, K. Maksymowicz, and M. Marciniak, “Mechanical properties of selective digestion of elastin and collagen from human aortas,” Acta of Bioengineering and Biomechanics; *02/2015; ISSN 1509-409X*, 2015, doi: 10.5277/ABB-00184-2014-02.

[12] M.-J. Chow, J. R. Mondonedo, V. M. Johnson, and Y. Zhang, “Progressive structural and biomechanical changes in elastin degraded aorta,” Biomech Model Mechanobiol, vol. 12, no. 2, pp. 361–372, Apr. 2013, doi: 10.1007/s10237-012-0404-9.

[13] S. Zeinali-Davarani, M.-J. Chow, R. Turcotte, and Y. Zhang, “Characterization of Biaxial Mechanical Behavior of Porcine Aorta under Gradual Elastin Degradation,” Ann Biomed Eng, vol. 41, no. 7, pp. 1528–1538, July 2013, doi: 10.1007/s10439-012-0733-y.

[14] M. Laffey, B. Tornifoglio, and C. Lally, “Development and Initial Characterisation of a Localised Elastin Degradation Ex Vivo Porcine Aortic Aneurysm Model,” Applied Sciences, vol. 13, no. 17, p. 9894, Sept. 2023, doi: 10.3390/app13179894.

[15] H. Weisbecker, C. Viertler, D. M. Pierce, and G. A. Holzapfel, “The role of elastin and collagen in the softening behavior of the human thoracic aortic media,” Journal of Biomechanics, vol. 46, no. 11, pp. 1859–1865, July 2013, doi: 10.1016/j.jbiomech.2013.04.025.

[16] K. Miura and K. Yamashita, “Mechanical Weakness of Thoracic Aorta Related to Aging or Dissection Predicted by Speed of Sound with Collagenase,” Ultrasound in Medicine & Biology, vol. 45, no. 12, pp. 3102–3115, Dec. 2019, doi: 10.1016/j.ultrasmedbio.2019.08.012.

[17] N. M. Ghadie, M. R. Labrosse, and J.-P. St-Pierre, “Glycosaminoglycans modulate compressive stiffness and circumferential residual stress in the porcine thoracic aorta,” Acta Biomaterialia, vol. 170, pp. 556–566, Oct. 2023, doi: 10.1016/j.actbio.2023.08.061.

[18] J. M. Mattson, R. Turcotte, and Y. Zhang, “Glycosaminoglycans contribute to extracellular matrix fiber recruitment and arterial wall mechanics,” Biomech Model Mechanobiol, vol. 16, no. 1, pp. 213–225, Feb. 2017, doi: 10.1007/s10237-016-0811-4.

[19] M.-J. Chow, M. Choi, S. H. Yun, and Y. Zhang, “The Effect of Static Stretch on Elastin Degradation in Arteries,” PLoS ONE, vol. 8, no. 12, p. e81951, Dec. 2013, doi: 10.1371/journal.pone.0081951.

[20] J. C. -Y. Chung et al., “Biomechanics of Aortic Dissection: A Comparison of Aortas Associated With Bicuspid and Tricuspid Aortic Valves,” JAHA, vol. 9, no. 15, p. e016715, Aug. 2020, doi: 10.1161/JAHA.120.016715.

[21] J. C.-Y. Chung et al., “Biomechanical properties of the aortic root are distinct from those of the ascending aorta in both normal and aneurysmal states,” JTCVS Open, vol. 16, pp. 38–47, Dec. 2023, doi: 10.1016/j.xjon.2023.08.015.

[22] M. Tang, D. Eliathamby, M. Ouzounian, C. A. Simmons, and J. C.-Y. Chung, “Dependency of energy loss on strain rate, strain magnitude and preload: Towards development of a novel biomarker for aortic aneurysm dissection risk,” Journal of the Mechanical Behavior of Biomedical Materials, vol. 124, p. 104736, Dec. 2021, doi: 10.1016/j.jmbbm.2021.104736.

[23] D. Eliathamby et al., “Biomechanics of the Aortic Root and Ascending Aorta in Patients With Marfan Syndrome,” European Journal of Cardio-Thoracic Surgery, vol. 67, no. 9, p. ezaf293, Sept. 2025, doi: 10.1093/ejcts/ezaf293.

[24] S. Peterson et al., “Regional differences in biomechanical properties of the ascending aorta in aneurysmal and normal aortas,” Biomech Model Mechanobiol, vol. 24, no. 3, pp. 865–877, June 2025, doi: 10.1007/s10237-025-01941-y.

[25] K. Cizkova, T. Foltynkova, M. Gachechiladze, and Z. Tauber, “Comparative Analysis of Immunohistochemical Staining Intensity Determined by Light Microscopy, ImageJ and QuPath in Placental Hofbauer Cells,” Acta Histochem. Cytochem., vol. 54, no. 1, pp. 21–29, Feb. 2021, doi: 10.1267/ahc.20-00032.

[26] B. Mirani et al., “Right Ventricular Stiffening and Function Are Associated With Main Pulmonary Artery Remodeling in a Rat Model of Pulmonary Hypertension,” ATVB, vol. 45, no. 6, pp. 945–964, June 2025, doi: 10.1161/ATVBAHA.124.321354.

[27] A. Cristoforetti, M. Masè, and F. Ravelli, “Model-Based Approach for the Semi-Automatic Analysis of Collagen Birefringence in Polarized Light Microscopy,” Applied Sciences, vol. 13, no. 5, p. 2916, Feb. 2023, doi: 10.3390/app13052916.

[28] P. Lacolley et al., “Disruption of the elastin gene in adult Williams syndrome is accompanied by a paradoxical reduction in arterial stiffness,” Clinical Science, vol. 103, no. 1, pp. 21–28, July 2002, doi: 10.1042/cs1030021.

[29] M. G. Jones et al., “Nanoscale dysregulation of collagen structure-function disrupts mechano-homeostasis and mediates pulmonary fibrosis,” eLife, vol. 7, p. e36354, July 2018, doi: 10.7554/eLife.36354.

[30] A. Gouissem, R. Mbarki, F. Al Khatib, and M. Adouni, “Multiscale Characterization of Type I Collagen Fibril Stress–Strain Behavior under Tensile Load: Analytical vs. MD Approaches,” Bioengineering, vol. 9, no. 5, p. 193, Apr. 2022, doi: 10.3390/bioengineering9050193.

[31] B. Depalle, Z. Qin, S. J. Shefelbine, and M. J. Buehler, “Influence of cross-link structure, density and mechanical properties in the mesoscale deformation mechanisms of collagen fibrils,” Journal of the Mechanical Behavior of Biomedical Materials, vol. 52, pp. 1–13, Dec. 2015, doi: 10.1016/j.jmbbm.2014.07.008.

[32] G. Fessel et al., “Advanced Glycation End-Products Reduce Collagen Molecular Sliding to Affect Collagen Fibril Damage Mechanisms but Not Stiffness,” PLoS ONE, vol. 9, no. 11, p. e110948, Nov. 2014, doi: 10.1371/journal.pone.0110948.

[33] D. Wågsäter et al., “Impaired Collagen Biosynthesis and Cross-linking in Aorta of Patients With Bicuspid Aortic Valve,” JAHA, vol. 2, no. 1, p. e000034, Jan. 2013, doi: 10.1161/JAHA.112.000034.

[34] J. E. Pichamuthu et al., “Differential Tensile Strength and Collagen Composition in Ascending Aortic Aneurysms by Aortic Valve Phenotype,” The Annals of Thoracic Surgery, vol. 96, no. 6, pp. 2147–2154, Dec. 2013, doi: 10.1016/j.athoracsur.2013.07.001.

[35] O. G. Andriotis, M. Nalbach, and P. J. Thurner, “Mechanics of isolated individual collagen fibrils,” Acta Biomaterialia, vol. 163, pp. 35–49, June 2023, doi: 10.1016/j.actbio.2022.12.008.

[36] C. A. Mariano, S. Sattari, G. O. Ramirez, and M. Eskandari, “Effects of tissue degradation by collagenase and elastase on the biaxial mechanics of porcine airways,” Respir Res, vol. 24, no. 1, p. 105, Apr. 2023, doi: 10.1186/s12931-023-02376-8.

[37] J. Z. Cui, A. Y. Tehrani, K. A. Jett, P. Bernatchez, C. Van Breemen, and M. Esfandiarei, “Quantification of aortic and cutaneous elastin and collagen morphology in Marfan syndrome by multiphoton microscopy,” Journal of Structural Biology, vol. 187, no. 3, pp. 242–253, Sept. 2014, doi: 10.1016/j.jsb.2014.07.003.

[38] J. Kim, M. C. Staiculescu, A. J. Cocciolone, H. Yanagisawa, R. P. Mecham, and J. E. Wagenseil, “Crosslinked elastic fibers are necessary for low energy loss in the ascending aorta,” Journal of Biomechanics, vol. 61, pp. 199–207, Aug. 2017, doi: 10.1016/j.jbiomech.2017.07.011.

[39] P. D. Weinberg, C. P. Winlove, and K. H. Parker, “The distribution of water in arterial elastin: Effects of mechanical stress, osmotic pressure, and temperature,” Biopolymers, vol. 35, no. 2, pp. 161–169, Feb. 1995, doi: 10.1002/bip.360350204.

[40] Y. Wang, J. Hahn, and Y. Zhang, “Mechanical Properties of Arterial Elastin With Water Loss,” Journal of Biomechanical Engineering, vol. 140, no. 4, p. 041012, Apr. 2018, doi: 10.1115/1.4038887.

[41] S. Roccabianca, G. A. Ateshian, and J. D. Humphrey, “Biomechanical roles of medial pooling of glycosaminoglycans in thoracic aortic dissection,” Biomech Model Mechanobiol, vol. 13, no. 1, pp. 13–25, Jan. 2014, doi: 10.1007/s10237-013-0482-3.

[42] R. K. June and D. P. Fyhrie, “Enzymatic digestion of articular cartilage results in viscoelasticity changes that are consistent with polymer dynamics mechanisms,” BioMed Eng OnLine, vol. 8, no. 1, p. 32, 2009, doi: 10.1186/1475-925X-8-32.

[43] D. J. Griffin, J. Vicari, M. R. Buckley, J. L. Silverberg, I. Cohen, and L. J. Bonassar, “Effects of enzymatic treatments on the depth-dependent viscoelastic shear properties of articular cartilage: MICROSCALE MECHANICS OF CARTILAGE DEGRADATION,” J. Orthop. Res., vol. 32, no. 12, pp. 1652–1657, Dec. 2014, doi: 10.1002/jor.22713.

[44] R. J. Van De Stadt, R. Kuijer, G. P. J. Van Kampen, M. H. M. T. De Koning, E. Van De Voorde-Vissers, and J. K. Van Der Korst, “Heterogeneity of proteoglycans extracted before and after collagenase treatment of human articular cartilage: I. Physical properties related to age,” Arthritis & Rheumatism, vol. 29, no. 10, pp. 1239–1247, Oct. 1986, doi: 10.1002/art.1780291009.

[45] R. Readioff, B. Geraghty, Y. A. Kharaz, A. Elsheikh, and E. Comerford, “Proteoglycans play a role in the viscoelastic behaviour of the canine cranial cruciate ligament,” Front. Bioeng. Biotechnol., vol. 10, p. 984224, Nov. 2022, doi: 10.3389/fbioe.2022.984224.

[46] M. Nightingale et al., “Biomechanics in ascending aortic aneurysms correlate with tissue composition and strength,” JTCVS Open, vol. 9, pp. 1–10, Mar. 2022, doi: 10.1016/j.xjon.2021.12.001.

[47] S.-I. Murtada and G. A. Holzapfel, “Investigating the role of smooth muscle cells in large elastic arteries: A finite element analysis,” Journal of Theoretical Biology, vol. 358, pp. 1–10, Oct. 2014, doi: 10.1016/j.jtbi.2014.04.028.

